# Fibroblast Stromal Support Model for Predicting Human Papillomavirus-Associated Cancer Drug Responses

**DOI:** 10.1101/2024.04.09.588680

**Authors:** Claire D. James, Rachel L. Lewis, Alexis L. Fakunmoju, Austin J. Witt, Aya H. Youssef, Xu Wang, Nabiha M. Rais, Apurva Tadimari Prabhakar, J. Mathew Machado, Raymonde Otoa, Molly L. Bristol

## Abstract

Currently, there are no specific antiviral therapeutic approaches targeting Human papillomaviruses (HPVs), which cause around 5% of all human cancers. Specific antiviral reagents are particularly needed for HPV-related oropharyngeal cancers (HPV^+^OPCs) whose incidence is increasing and for which there are no early diagnostic tools available. We and others have demonstrated that the estrogen receptor alpha (ERα) is overexpressed in HPV^+^OPCs, compared to HPV-negative cancers in this region, and that these elevated levels are associated with an improved disease outcome. Utilizing this HPV^+^ specific overexpression profile, we previously demonstrated that estrogen attenuates the growth and cell viability of HPV^+^ keratinocytes and HPV^+^ cancer cells *in vitro*. Expansion of this work *in vivo* failed to replicate this sensitization. The role of stromal support from the tumor microenvironment (TME) has previously been tied to both the HPV lifecycle and *in vivo* therapeutic responses. Our investigations revealed that *in vitro* co-culture with fibroblasts attenuated HPV^+^ specific estrogen growth responses. Continuing to monopolize on the HPV^+^ specific overexpression of ERα, our co-culture models then assessed the suitability of the selective estrogen receptor modulators (SERMs), raloxifene and tamoxifen, and showed growth attenuation in a variety of our models to one or both of these drugs *in vitro.* Utilization of these SERMs *in vivo* closely resembled the sensitization predicted by our co-culture models. Therefore, the *in vitro* fibroblast co-culture model better predicts *in vivo* responses. We propose that utilization of our co-culture *in vitro* model can accelerate cancer therapeutic drug discovery.

**Importance:** Human papillomavirus-related cancers (HPV^+^ cancers) remain a significant public health concern, and specific clinical approaches are desperately needed. In translating drug response data from *in vitro* to *in vivo*, the fibroblasts of the adjacent stromal support network play a key role. Our study presents the utilization of a fibroblast 2D co-culture system to better predict translational drug assessments for HPV^+^ cancers. We also suggest that this co-culture system should be considered for other translational approaches. Predicting even a portion of treatment paradigms that may fail *in vivo* with a co-culture model will yield significant time, effort, resource, and cost efficiencies.

## Introduction

Human papillomaviruses (HPVs) are small, double-stranded DNA viruses, and high-risk HPVs are known carcinogens^1–7^. HPV is the most common sexually transmitted infection in the United States (U.S.), and estimated to infect more than 80% of the population at least once in their lifetime^1–3,8–12^. HPV16 is the most prevalent genotype, accounting for at least 50% of cervical cancers and approximately 90% of HPV^+^ oropharyngeal cancers (HPV^+^OPCs)^9,13,14^. While prophylactic HPV vaccines have already begun to show remarkable efficacy in preventing infection and related diseases, HPV continues to account for ∼5% of worldwide cancer, and disproportionately affects marginalized populations both in the U.S. and around the world^1–3,8–12,15–17^. As such, the lack of specific antiviral therapeutics available for combatting HPV-related cancers is of significant concern.

While many cancers are on the decline, the last two decades have shown a sharp increase in HPV^+^OPCs, for which there are no early diagnostic tools available^4–6,18,19^. HPV^+^OPCs are found at 4-fold higher levels in men than in women, suggesting there are sex-related differences in the development of these cancers^4,18^. Using data from The Cancer Genome Atlas (TCGA), we and others have shown that the estrogen receptor alpha (ERα) is overexpressed in HPV^+^HNC (head and neck cancers including OPCs) and that these elevated levels are associated with an improved disease outcome^18,20–26^.

We have also previously demonstrated that 17-β estradiol (estrogen) attenuates the growth and cell viability of HPV^+^ keratinocytes and HPV^+^ cancer cells *in vitro*, but not HPV negative (HPV^-^) keratinocytes or HPV^-^ cancer cells^26^. Sensitization occurs via numerous mechanisms: 1) at the level of viral transcription, 2) via interactions with E6 and E7, 3) through manipulation of cell survival and cell death pathways^26^. Here, we report that the expansion of estrogen treatment into *in vivo NOD-scid IL2Rg^null^* (NSG) mice revealed a lack of response to estrogen alone or in combination with radiotherapy (IR).

Previously, estrogen has been shown to promote HPV-induced cervical disease in immunocompetent mice, yet this enhancement was lost in NSGs^27–32^. HPV oncogenes, in conjunction with estrogen, were shown to fundamentally reprogram the tumor microenvironment (TME)^27,28,33^. We therefore sought to determine if the TME, specifically the stromal support provided by fibroblasts, alters the estrogenic effects in our model systems^33–42^. *In vitro* co-culture studies revealed that stromal interactions markedly change cell growth and viability in response to estrogen in our HPV^+^ models.

While our estrogen studies did not prove effective *in vivo*, ERα remains overexpressed in HPV^+^OPC and HPV^+^ cervical cancers^24–27,29,31–33,43^. Selective estrogen receptor modulators (SERMs) have proven to provide a multitude of therapeutic applications^44–50^. Analysis of K14E6/E7 transgenic mice models have previously shown the possible utility of raloxifene on reduction of recurrence of cervical neoplastic disease, and earlier literature suggested the utility of tamoxifen to lengthen the latent period for cervical dysplasia and carcinoma in carcinogen-induced models^45,51^. We utilized our co-culture system to assess the efficacy of raloxifene and tamoxifen and showed significant growth repression to one or both SERMs in a number of cancer cell lines. Our preliminary assessment of these SERMs in an *in vivo* HPV^+^HNC cell line correlated with our *in vitro* observations and suggests that these drugs may be useful adjuvant approaches for further investigation. To our knowledge, our analysis is the first to provide evidence on the utility of SERMs for HPV^+^OPCs.

Overall, this report further expands upon the analysis of utilizing estrogen related signaling in the quest for HPV-specific antiviral approaches^25^. While estrogen presented compelling evidence *in vitro*, this report demonstrates that estrogen treatment did not translate to *in vivo* models^26^. The development of co-culture models utilizing fibroblast “feeder” cells demonstrated that these supporting fibroblasts altered the response to estrogen *in vitro*, modeling what was observed *in vivo*. Analysis of SERMs with this co-culture model demonstrated the utility of altering estrogen-related signaling both *in vitro* and *in vivo*. Therefore, co-culture with fibroblasts offers a simple and more physiologically relevant environment by stimulating more of the cellular interactions present in solid tumors. This co-culture model may serve to better predict drug responses in other translational paradigms, not limited to HPV^+^ cancers. Co-culture allows for an examination of the complex cellular responses to drug effects *in vivo*, thereby enhancing the accuracy of *in vitro* therapeutic evaluations for more successful translational approaches.

## Results

### Estrogen sensitization is not observed *in vivo*

We have previously reported that estrogen attenuates the growth of epithelial cells in an HPV^+^ dependent manner *in vitro*^26^. We found that this occurred via both a repression of transcription from the HPV16 long control region and through interactions with the viral oncogenes, E6 and E7^26^. Furthermore, estrogen treatment enhanced irradiation-induced cell death in an HPV^+^ dependent manner^26^. A logical progression was to assess the response of HPV^+^ cancers to estrogen *in vivo*. Previously, our laboratory as well as others have demonstrated that HeLa cells are highly responsive to estrogen treatment^26,52,53^. Consequently, experiments were designed to assess the combination of estrogen and radiation treatment on HeLa xenografts in female NSG mice. Contrary to our *in vitro* data, Figure 1A shows that estrogen alone, or in combination with radiation, had no impact on tumor response in mice. Of note, animal weights remained consistent throughout the study with all treatments (Figure 1B).

**Figure 1:**
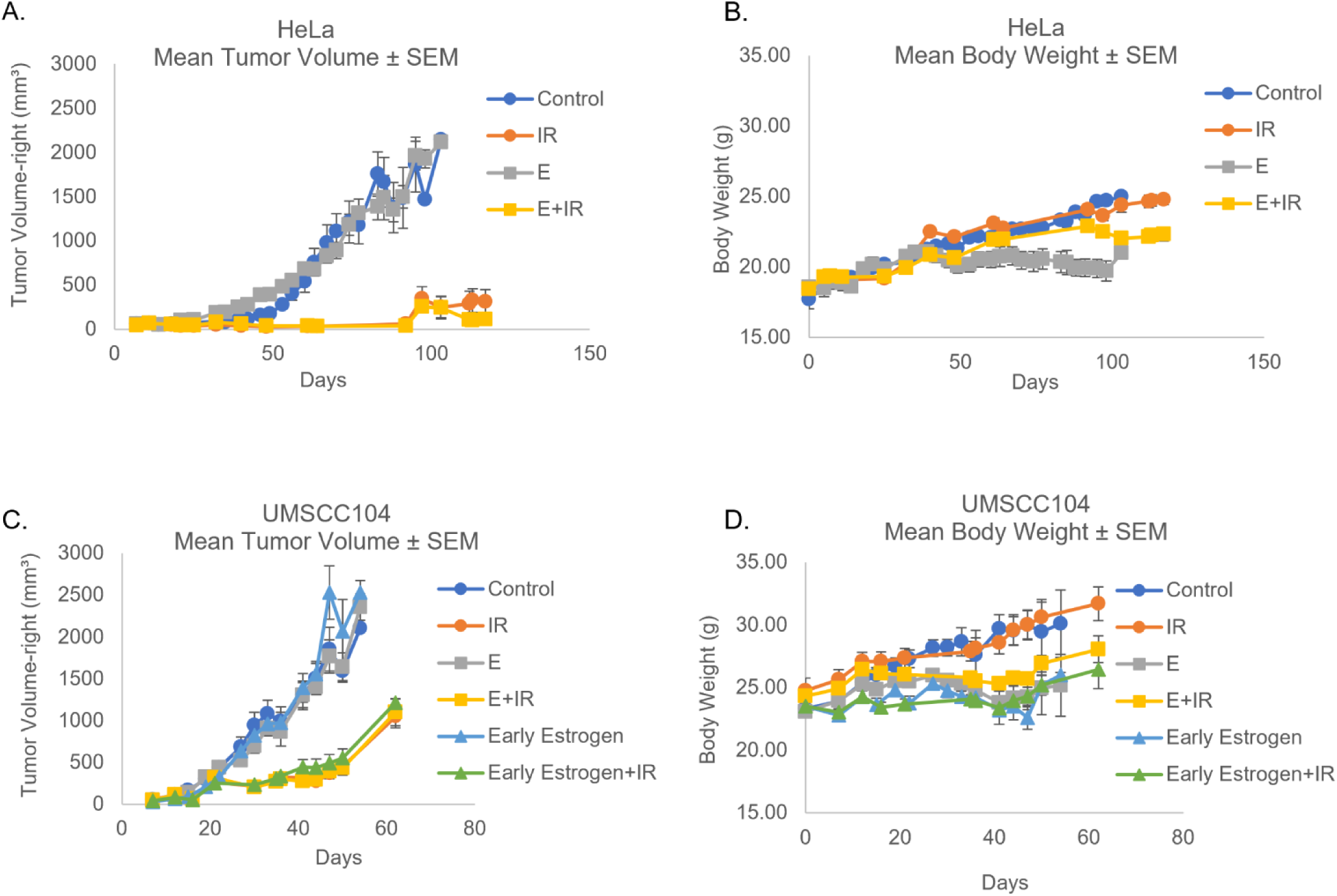
Estrogen fails to sensitize *in vivo*. 1A. HeLa cells are an integrated HPV18+ female cervical cancer cell line, we therefore chose to utilize female NOD-scid IL2Rgnull (NSG) mice for this treatment paradigm; we did not choose to ovariectomize these mice. Day 0 marks the date at which cells were injected for xenografts. Tumors were palpable on day 10, mice were randomized. Estrogen alone (E=0.4mg pellet + 8 µg/ml water supplementation *ad libitum*), radiation alone (IR=10 Gy), as well as the combinational approach (E+IR) were monitored for effects on tumor volume by calipers. 1B. Mice were monitored for weight throughout the study. 1C. UMSCC104 cells are an episomal HPV16+ male oropharyngeal line, we therefore chose to utilize male NSG mice for this treatment paradigm. Day 0 marks the date at which cells were injected for xenografts; early estrogen water supplementation began on this day, pellets were later added when tumors were palpable (8 µg/ml water supplementation *ad libitum*). Tumors were palpable on day 7. Estrogen alone (0.4mg pellet + 8 µg/ml water supplementation ad libitum), radiation alone (5 Gy), as well as the combinational approach were monitored for effects on tumor volume by calipers. 1D. Mice were monitored for weight throughout the study.

Previously, we had observed HPV^+^ dependent cell death following estrogen treatment regardless of sex, tissue of origin, or viral genome integration status^26^. To determine if any of these factors played a role *in vivo*, we decided to expand our analysis to include the male, episomal, HPV16^+^ oropharyngeal cancer cell line UMSCC104^54^. In addition, we added “early” estrogen supplementation that began the same day as xenografts were injected, to determine if a temporal relationship was essential to the estrogen treatment response. As our previous radiation treatment using 10 Gy had some off-target effects in our NSGs, radiation was reduced to 5 Gy in these studies. As observed in Figure 1C (animal weight Figure 1D), estrogen again had no impact on tumor response in any of these conditions. These results complement the previous observations by the Lambert laboratory utilizing MmuPV1^27^. The Lambert observations demonstrated that estrogenic effects *in vivo* are reliant, at least in part, on estrogen’s suppression of the host immune system in immunocompetent mice; whereas estrogenic enhancement of disease progression were not observed in immunodeficient NSG mice^27^. Our results indicate that the lack of estrogenic alterations in disease progression may not be papillomavirus species specific, but further studies are needed to confirm this observation.

### Stroma alters HPV-specific estrogen growth response in keratinocyte models

There are numerous differences when moving from *in vitro* to *in vivo* models. An increasingly recognized component of *in vivo* responses is the adjacent stromal support network, or “tumor microenvironment” (TME)^33,55,56^. In regard to HPV, evidence also supports a significant role for stroma during the viral lifecycle and HPV-induced disease ^33,39,41,42^. Fibroblasts are a key component of this stromal support and can significantly alter cancer resistance and therapeutic responses^33–42^. We already utilize feeder layers of mitomycin C (MMC) growth-arrested murine 3T3-J2 fibroblasts (referred to as J2s moving forward) in the immortalization and maintenance of our primary keratinocyte derived HPV^+^ epithelial cultures. Thus, we applied the same approach to investigate whether this co-culture system affects the response to estrogen. MMC inactivation is a supported approach to arrest the proliferation of fibroblasts, allowing for the establishment of a supportive feeder layer that maintains its ability to synthesize RNA and protein and provide stromal regulation of neighboring cells of interest^57^. This approach is widely accepted as necessary for maintenance of the HPV genome, as well as a necessary component of 3D models for HPV lifecycle analysis^58–64^. N/Tert-1 cells (telomerase immortalized foreskin keratinocytes, HPV negative), as well as HFK+E6E7 (foreskin keratinocytes immortalized by the viral oncogenes only), and HFK^+^HPV16 (foreskin keratinocytes immortalized by the entire HPV16 genome, replicating as an episome), were cultured in the presence or absence of J2 cells, and treated with 15µM estrogen, or vehicle control^26^. Compared to untreated, non-cocultured N/Tert-1 cells, no significant alterations in growth rate were observed following estrogen treatment or co-culture with J2s (Figure 2A). Conversely, estrogen significantly repressed the growth of both the E6E7 and HPV16 immortalized cell lines in the absence of stromal support (Figure 2B and 2C, respectively). The presence of stromal support fibroblasts significantly rescued this growth suppression (Figure 2B and 2C); coculture with fibroblasts mitigates the growth suppressive effects of estrogen, *in vitro*. This implies that the presence of stromal support is responsible for the differences in response to estrogen treatment that we observed between our *in vitro* and *in vivo* models (Figure 1)^26^.

**Figure 2:**
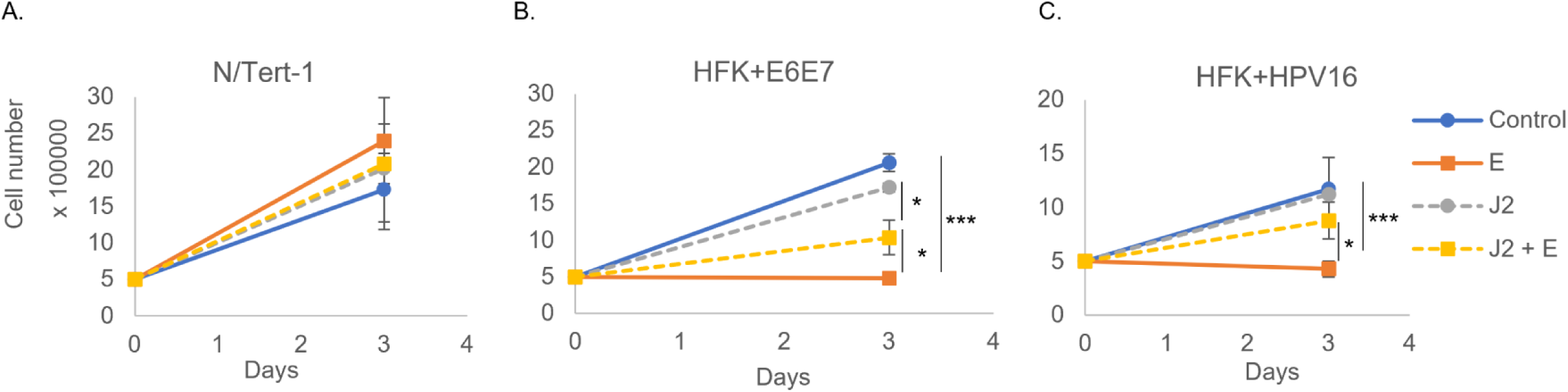
Fibroblasts significantly reduce HPV-specific estrogenic sensitization in keratinocytes. (A) N/Tert-1, (B) HFK+E6E7, and (C) HFK+HPV16 cells were seeded on day 0 and grown in the presence or absence of J2s that had been seeded at least 6 hours prior. Cells were washed to remove J2s in noted conditions, trypsinized, and counted on day 3. *, p < 0.05; ***, p < 0.001.

### Stroma alters HPV-specific estrogen growth response in cancer models

We sought to expand our co-culture investigations to include cancer lines. A hallmark of transformation in many cancer cell lines is anchorage independent growth and loss of adherence in cell culture^65^. To improve our cancer cell count analysis, we developed a novel, quantifiable co-culture system. Using the Nuclight lentivirus system (Sartorius), we transduced our cells of interest with nuclear mKate2-red and developed stable cell lines; this system enabled automated cell counting and could distinguish between our cells of interest and the non-labeled J2s. An additional advantage of this system was the ability to monitor cellular morphology and observe any co-culture influences upon colony formation and cellular distribution. We sought to determine whether J2s alter cancer cell line responses to estrogen, as observed in our keratinocyte models (Figure 2). We utilized four cancer cell lines in our co-culture experiments: HN30 – an HPV^-^ head and neck cancer (p53wt), HeLa – an HPV18^+^ integrated cervical cancer (p53wt), UMSCC47 – an HPV16^+^ integrated head and neck cancer (p53wt), and UMSCC104 – an HPV16^+^ episomal head and neck cancer (p53wt). As HeLa cells are highly sensitive to estrogen, 1.5µM estrogen was utilized in experiments with these cells; all other cell lines were treated with 15µM as previously described^26,52^. As previously shown, estrogen treatment significantly repressed cell growth in an HPV^+^ dependent manner (Figures 3A,C,E,F)^26^. Conversely, estrogenic sensitivity was no longer observed in HPV^+^ cancer cells grown with feeder cells (Figures 3B,D,F,H). This was similar to the loss of estrogenic sensitivity in our HPV^+^ keratinocytes (Figure 2), and in our mouse studies (Figure 1A, and 1C). Again, demonstrating that stromal support alters estrogenic sensitivity of HPV^+^ cancer models.

**Figure 3:**
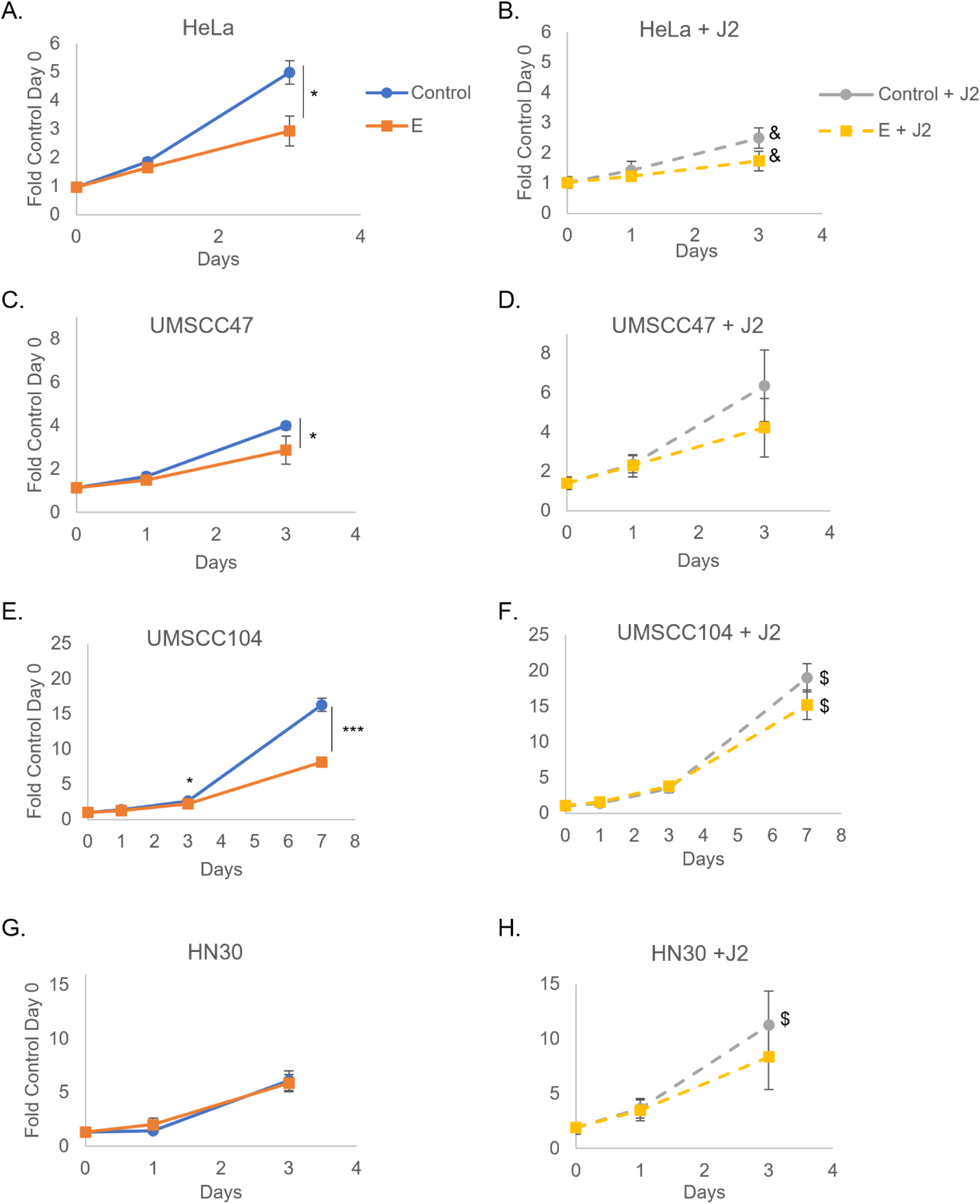
Fibroblasts significantly reduce HPV-specific cancer estrogenic sensitization. J2s were seeded in the morning and noted nuclear-labeled cancer cells were seeded at least 6 hours after: HeLa (A,B), UMSCC47 (C,D), UMSCC104 (E,F), HN30 (G,H). Co-culture images for quantitation were taken the following morning and are set at day 0, estrogen (E) was added immediately after initial imaging on day 0. Cells were again imaged at day 1 and day 3. UMSCC104 cells were grown for an additional time point; these were replenished with new J2s and estrogen on day 3 (post-imaging) and day 5 and imaged again on day 7. Within same graphs *p<0.05, ***p<0.001. J2s altered growth rates for some of the cell lines and graphs are presented as separate for visual simplicity, but experiments were run concurrently; comparing top and bottom graphs $p<0.05 J2 increased growth, &p<0.05 J2 decreased growth.

To determine if this repression of estrogenic sensitization was specific to mouse fibroblast support in co-culture, HeLa cells were grown in the presence or absence of conditioned media collected from replicating 3T3-J2s cultures, or grown in co-culture with MMC-inactivated human dermal mesenchymal fibroblasts (HDFMs). Again Figure 4A demonstrates that HeLa cells in monoculture undergo significant growth repression in the presence of estrogen. Similarly, estrogenic growth repression was observed in the presence of J2 conditioned media (Figure 4B). Co-culture with HDFMs rescued estrogenic growth repression in HeLa cells (Figure 4C), similar to what was observed with J2s (Figure 3B). HDFMs did not repress the growth of HeLa alone (Figure 4C), contrary to the growth repression observed in the presence with J2 (Figure 3B). While we do not currently understand the mechanism behind this altered growth potential with alternative species’ fibroblasts, we find it noteworthy that estrogenic sensitivity is lost with both co-culture methods, but not by incubation with media, indicating that cell-cell contact is required. Moreover, this loss of sensitivity is not dependent on fibroblast growth alteration of cancer cells.

**Figure 4:**
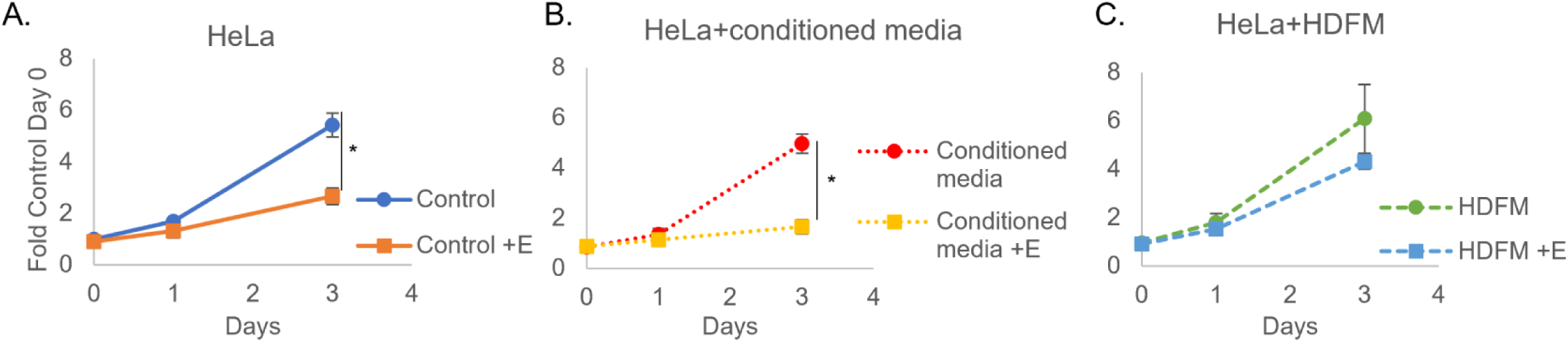
Human fibroblasts (HDFM) significantly reduce HPV-specific cancer estrogenic sensitization. HeLa were seeded into standard media (A), J2 conditioned media (B), or HDFMs were seeded in the morning and nuclear-labeled HeLa cells were seeded at least 6 hours after (C). Co-culture images for quantitation were taken the following morning and are set at day 0, estrogen (E) was added immediately after initial imaging on day 0. Cells were again imaged at day 1 and day 3. \**p*<0.05

### Stroma does not alter the response to cisplatin in cancer models

To assess the predictiveness of our co-culture model, we found it important to investigate whether stroma would alter the response to other chemotherapeutic approaches. HeLa, UMSCC47, UMSCC104, and HN30 cells were therefore evaluated for their responsiveness to cisplatin (Figure 5A,C,E,G). Fibroblasts did not rescue the growth arrest observed in all cell lines (Figures 5B,D,F,H). Cisplatin is a well-established clinical treatment paradigm for HPV^+^ and HPV^-^OPCs, cervical cancer, and many other cancers^66–74^. We predicted that fibroblasts would not be able to rescue cancer cells from this conventional chemotherapeutic agent. We postulate that fibroblasts are unlikely to change the response to most currently accepted cancer treatment modalities. Instead, we suggest that this model may better predict which novel therapeutics may fail translational approaches. This is something we are currently investigating, and we encourage others to consider this approach as well when translating treatments from *in vitro* to *in vivo*.

**Figure 5:**
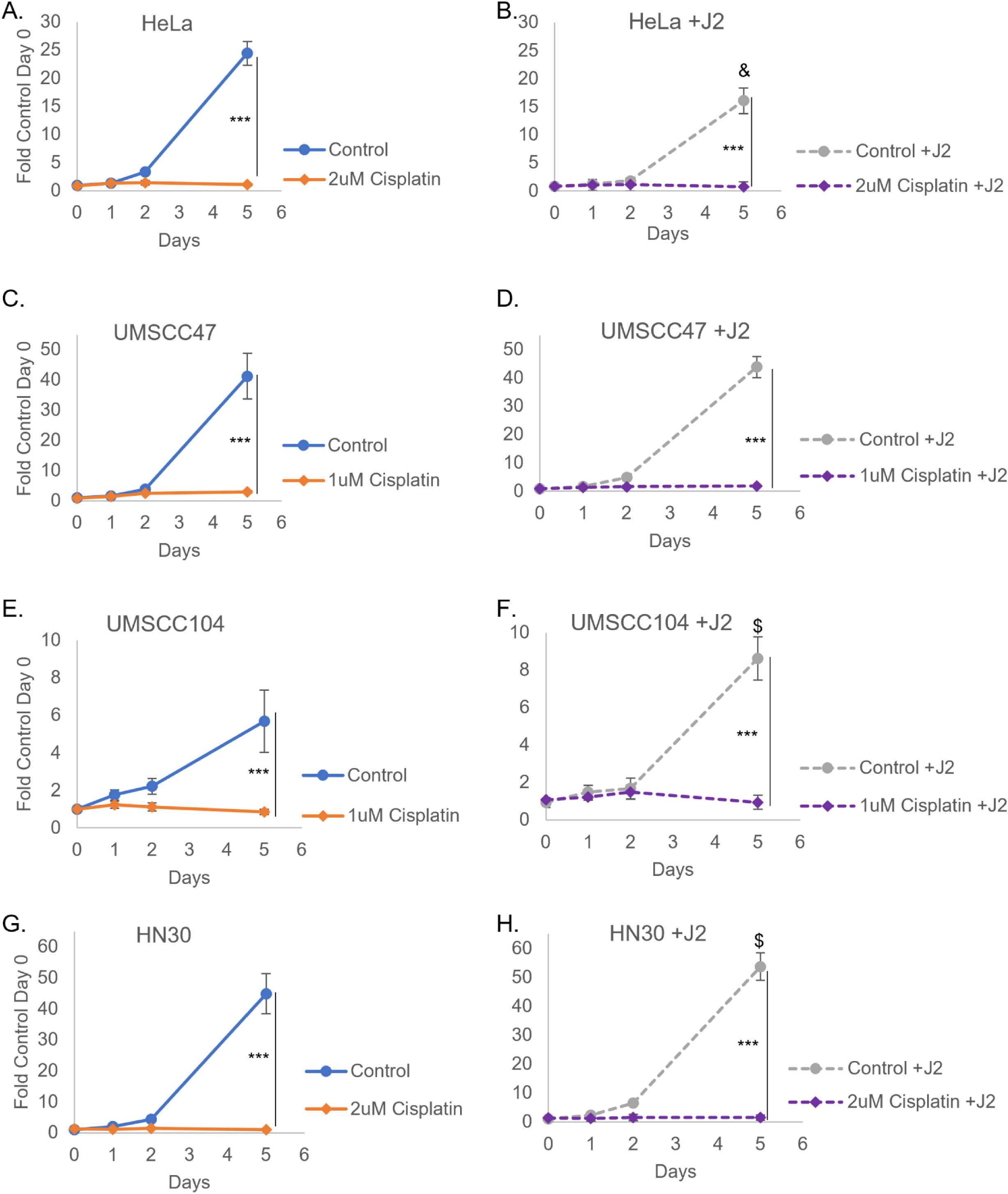
Fibroblasts do not alter cisplatin sensitivity. J2s were seeded in the morning and noted nuclear-labeled cancer cells were seeded at least 6 hours after: HeLa (A,B), UMSCC47 (C,D), UMSCC104 (E,F), HN30 (G,H). Co-culture images for quantitation were taken the following morning and are set at day 0, noted concentrations of cisplatin were added immediately after initial imaging on day 0. Cells were again imaged at day 1 and day 2. New J2s or media were replenished on day 3, and imaged again on day 5. Within same graphs \*\*\*\**p*<0.001. J2s altered growth rates for some of the cell lines and graphs are presented as separate for visual simplicity, but experiments were run concurrently; comparing top and bottom graphs $*p*<0.05 J2 increased growth, &*p*<0.05 J2 decreased growth.

### Stroma alters HPV long control region (LCR) transcriptional regulation

We previously published that estrogen represses transcription of the HPV16 long control region (LCR) both in our N/Tert-1 and C33a transcription models^26^. This transcriptional repression of the LCR, in turn, downregulated the expression of early viral genes in our numerous HPV^+^ keratinocyte and HPV^+^ cancer models^26^. Expanding upon these previous observations, we further wanted to determine the impact of stromal support on estrogenic HPV16-LCR transcriptional regulation. N/Tert-1 cells grown in the presence of stromal support had significantly enhanced levels of HPV16-LCR transcription and this transcriptional regulation was no longer significantly repressed by estrogen (Figure 6A); this may be one of the many contributing factors involved in the loss of estrogen sensitivity. ERα was also assessed, and stroma did not appear to alter protein expression in N/Tert-1 cells (Figure 6B). These results suggest that stroma is highly supportive of HPV16-LCR transcriptional regulation.

**Figure 6:**
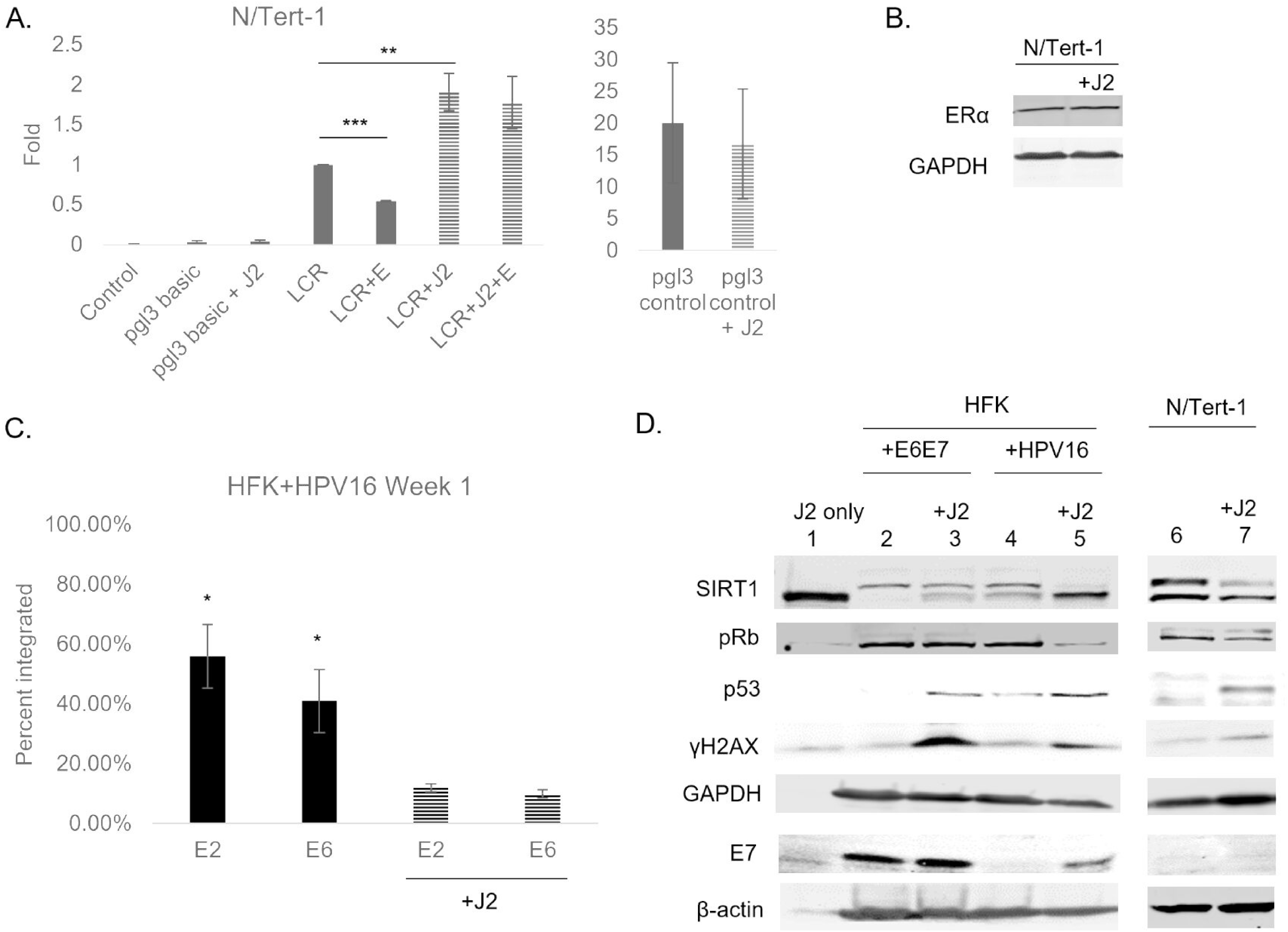
Stroma supports transcriptional regulation, viral protein expression, and episomal maintenance; estrogenic transcriptional regulation is lost with stroma. (A) N/Tert-1 cells were transfected with 1 μg of pgl3 basic backbone, 1 μg of pgl3 control (positive control), or 1 μg LCR and grown in the presence or absence of 15 μM estrogen and/or J2s that had previously been seeded. Forty-eight hours after transfection, a luciferase-based assay was utilized to monitor levels of LCR transcription. Data are presented as relative fluorescence units (RFU), normalized to total protein concentration as monitored by a standard bovine serum albumin (BSA) assay. ANOVA **, *P* < 0.01; ***, *P* < 0.001 (B) N/Tert-1 cells were grown in the presence or absence of J2s. Cells were washed to remove J2, then lysed and analyzed via western blotting for ERα. GAPDH was used as a loading control. (C) HFK^+^HPV16 cells were seeded on day 0 and grown in the presence or absence of J2s for 1 week. Cells were washed to removed J2, then lysed and analyzed for DNA expression of E2 and E6 via the exonuclease V assay, in comparison to GAPDH and mitochondrial DNA controls. Results are presented as percent integration as calculated from the cut ratio of matched GAPDH. \**P <* 0.05. (D) HFK+E6E7 (lanes 2,3) and HFK^+^HPV16 (lanes 4,5) and N/Tert-1 (lanes 6,7) cells were seeded on day 0 and grown in the presence or absence of J2s for 1 week. Lane 1 is an input lysate from J2 alone to control for any background level of expression in fibroblasts that were not removed via washing. Cells were washed to remove J2s in noted conditions, trypsinized, lysed, and analyzed via western blotting for SIRT1, pRb, p53, γH2aX, and E7. GAPDH and β-actin were utilized as loading controls.

Fibroblasts are routinely utilized to support HPV genome maintenance and the viral life cycle in keratinocyte models^33,41,57,75–82^. While this model is accepted, the full mechanism of how fibroblasts aid maintenance of the viral genome as an episome has yet to be elucidated. Figure 6A demonstrates that stromal support enhances HPV16-LCR transcription. It was therefore important to confirm that this transcriptional regulation had downstream effects. Previously, elements of the HPV upstream regulatory region, another term for the viral LCR, have been implicated as a requirement for long term viral persistence in keratinocytes^83^. The McBride laboratory demonstrated that the chromatin architecture of this region is important for genome partitioning and may influence integration^83^. Figure 6C confirms that HFK^+^HPV16 begin to integrate the viral genome when grown in the absence of J2 for one week^84,85^. The accepted practice of fibroblast co-culture, may in part maintain viral episomes via influencing the transcription of LCR. Thus, stroma supports viral protein expression and alters host signaling pathways in keratinocytes

We next investigated viral protein expression and host protein signaling observed in the presence or absence of fibroblast support. HFK+E6/E7, HFK^+^HPV16, and N/Tert-1 cells (Figure 6D) were grown in the presence or absence of J2s. J2s are washed off before harvesting samples, and are murine-derived so there should be limited detection via human-specific antibodies. To confirm that altered protein levels observed were not due to any residual J2s, lane 1 in Figure 6D demonstrates that significant specific bands were not observable with the majority of our chosen antibodies with 100µg of J2 protein input. SIRT1 does have a notable lower band which is not unexpected due to the gene homology between mouse and human SIRT1 and that the immunogen was developed from the amino acids 1-131 of mouse Sir2α. γH2AX has a low-level detectible band due to cross reactivity of the antibody; it should be noted that the significant observations discussed have taken this into account. Figure 6D demonstrates that HFK+E6E7, which do not rely on the LCR for early gene expression, do not have altered E7 protein levels in the presence or absence of fibroblasts (lanes 2,3). As fibroblasts were shown to enhance LCR transcription (Figure 6A), HFK^+^HPV16 cells, which rely on the LCR for early gene expression, have enhanced E7 levels when grown in the presence of J2s (lanes 4,5). HFK+E6E7 had limited levels of p53 expression, and HFK^+^HPV16 had low expression of p53, due to E6-targeted degradation; however significant enhancement of p53 levels in the presence of J2 in both cell lines were striking^63,64,79,86–91^. To our knowledge, this is the first report to suggest why p53 is not always fully degraded in HPV^+^ cell lines; E6 degradation of p53 may depend on whether or not keratinocytes are maintained on feeder cells^79,88,90,92–97^. Upregulation of p53 in the presence of J2 was also observed in N/Tert-1 cells (lanes 6,7) so this observation is independent from E6 or full genome expression. It is worth noting that HFK^+^HPV16 cells are consistently maintained in co-culture with J2s, and that J2s were not supplemented for the “control” conditions. This also demonstrates that the stromal induced alterations of p53 protein levels are reversible (Figure 6D). Total levels of pRb were not altered N/Tert-1 cells nor in HFK+E6E7; this corresponds to unaltered E7 levels in HFK+E6E7 cells. Alternatively, pRb was reduced in HFK^+^HPV16 grown in the presence of J2s. Again, suggesting fibroblasts are important for viral regulation of keratinocyte signaling, possibly through the LCR. A consistent observation was the upregulation of γH2AX in the presence of J2s in all of the cell lines (Figure 6D). Numerous reports have demonstrated HPV viral integrity and genome stability is highly reliant on DNA repair machinery, including γH2AX; J2 enhancement of γH2AX may be another key mechanism of fibroblast regulation of viral genome stability in keratinocyte models^7,76,98–106^. Upstream of p53 and γH2AX, SIRT1 is known to target histones and non-histone substrates such as p53, and has been shown to decrease in response to DNA damage^107^. N/Tert-1 and HFK^+^HPV16 grown in the presence of J2s demonstrated a significant decrease in SIRT1. The lower band observed can be associated with SIRT1 post-translational modifications; while this was altered by fibroblasts in both HFK+E6E7 and HFK^+^HPV16, conclusions were not made due to the band observed with the J2 input sample ^40,107–110^. While we are still investigating additional signaling mechanisms involved, these observations highlight the importance of properly controlling for whether or not cell lines are grown in the presence or absence of feeder cells when considering viral and host signaling events.

### Selective estrogen receptor modulators reduce growth rates *in vitro* and *in vivo*

While estrogen had no impact on the tumor response in mice, alone or in combination with radiation (Figure 1), we expanded our analysis to assess if the selective estrogen receptor modulators (SERMs) raloxifene or tamoxifen would be a useful approach to continue to take advantage of the HPV^+^ specific overexpression of ERα. It should be noted that SERMs can be agonists or antagonists of the ERα, and these responses are dependent on region, cell type, and the localization and availability of estrogen response elements (EREs)^44,46,47,111^. Preclinical data supports the utility of SERMs for cervical cancer, particularly the utility of raloxifene on the reduction of recurrent neoplastic disease; however, SERMs have yet to be evaluated in HPV^+^OPC^47,111^.

HeLa, UMSCC47, and UMSCC104 HPV^+^ cancer cells grown without fibroblast support exhibited significant growth repression to both SERMs (Figures 7A,C,E). While fibroblast support did not alter the response to SERMs in all the cell lines, rescued growth was observed in HeLa cells treated with tamoxifen while remaining responsive to raloxifene (Figures 7B,D,F). It is worth noting that previous *in vivo* and clinical analysis have predicted the utility of raloxifene in HPV^+^ cervical cancer response, whereas tamoxifen is not recommended^45–47,51,111^. We therefore sought to analyze SERM responsiveness in an HPV^+^HNC *in vivo* model. Expansion of SERM treatment *in vivo* was therefore conducted in UMSCC104 cells alone or in combination with IR to assess their utility as well as determine if our co-culture model could be useful in future translational approaches. For this study, 4 Gy IR was chosen to further reduce the off-target issues relating to radiation use in NSGs^112^. As observed in Figure 8A, radiation alone significantly reduced tumor volume starting on Day 14, while raloxifene (Figure 8A) and tamoxifen alone (Figure 8B) were able to significantly reduce tumor volume starting on Day 32. Furthermore, in comparison to radiation (IR) alone, tamoxifen+IR significantly reduced tumor volume starting on Day 42 (Figure 8B). Kaplan-Meier survival analysis demonstrated 50% survival for tamoxifen+IR on day 70 in comparison to 10% survival at the same time point for IR alone or raloxifene+IR at our chosen endpoint (Figure 8C). With all treatments, animal weights remained consistent throughout the study (Figure 8D). Our *in vitro* co-culture system modeled our *in vivo* responses, as well as those previously observed, to both estrogen and SERMs in HPV^+^ cervical *in vivo* models (Figure 1,3,7) ^27,32,43,45–47,51,111^. Co-culture also predicted the responsiveness of SERMs in an HPV^+^HNC *in vivo* model (Figure 7,8) and suggests that the utility of SERMs for the treatment of HPV^+^OPC are worth further investigation. Altogether, we suggest that the predictiveness of this co-culture system should be considered in more translational approaches. Current studies are investigating the molecular mechanisms behind the alterations observed when cells are grown in fibroblast co-culture. These mechanistic approaches may further expand the predictive utility of our co-culture model.

**Figure 7:**
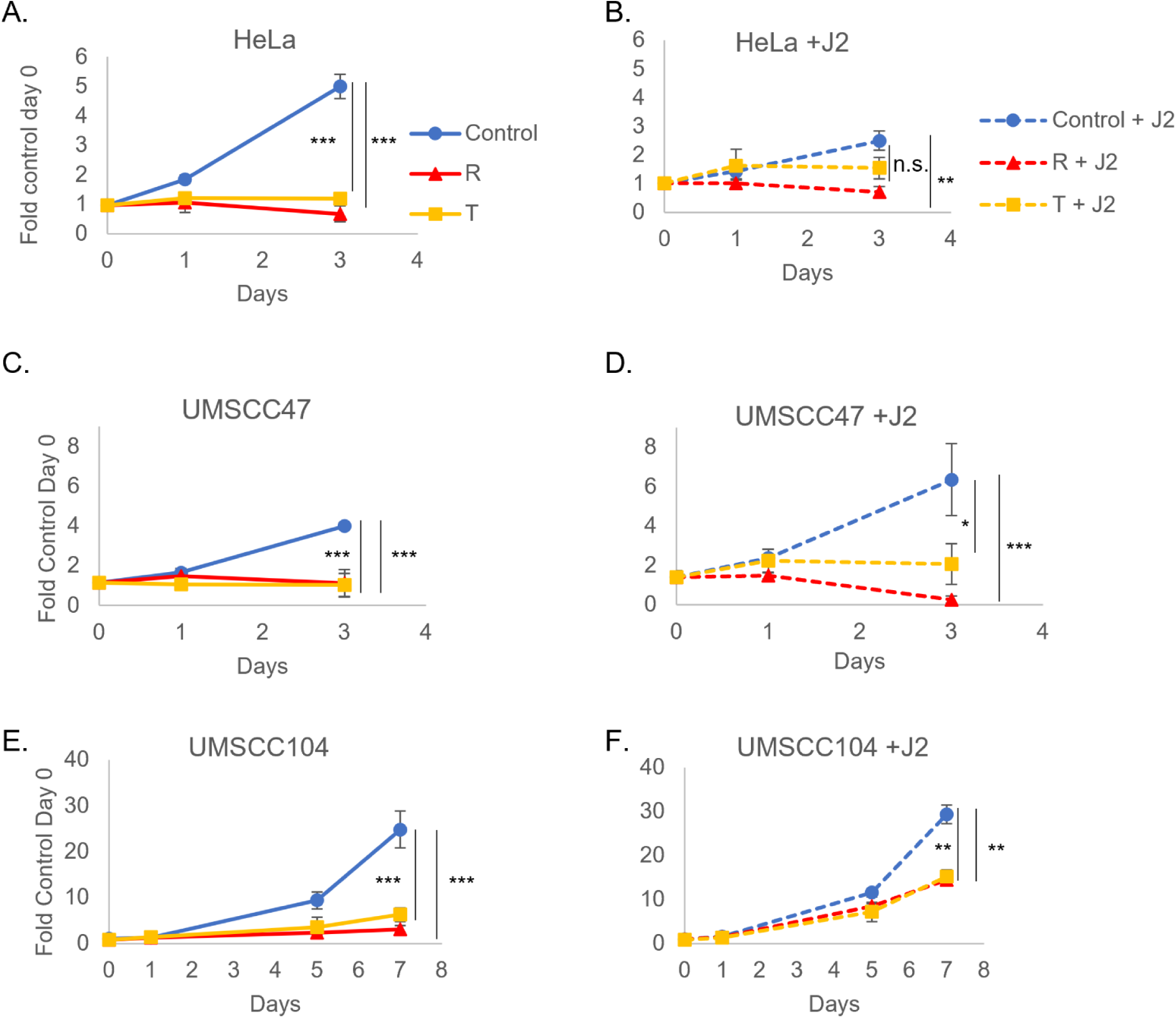
Fibroblast co-culture demonstrates SERMs are worth assessing in an HPV+ *in vivo* model. J2s were seeded in the morning and noted nuclear-labeled cancer cells were seeded at least 6 hours after: HeLa (A,B), UMSCC47 (C,D), UMSCC104 (E,F). Co-culture images for quantitation were taken the following morning and are set at day 0, 10 µg raloxifene (R) or 10 µg tamoxifen (T) were added immediately after initial imaging on day 0. Cells were again imaged at day 1 and day 3. UMSCC104 cells were grown for an additional time point; these were replenished with new J2s and drugs on day 3 (post-imaging) and day 5 and imaged again on day 7. Within same graphs *p<0.05, ***p<0.001. J2s altered growth rates for some of the cell lines and graphs are presented as separate for visual simplicity, but experiments were run concurrently.

**Figure 8:**
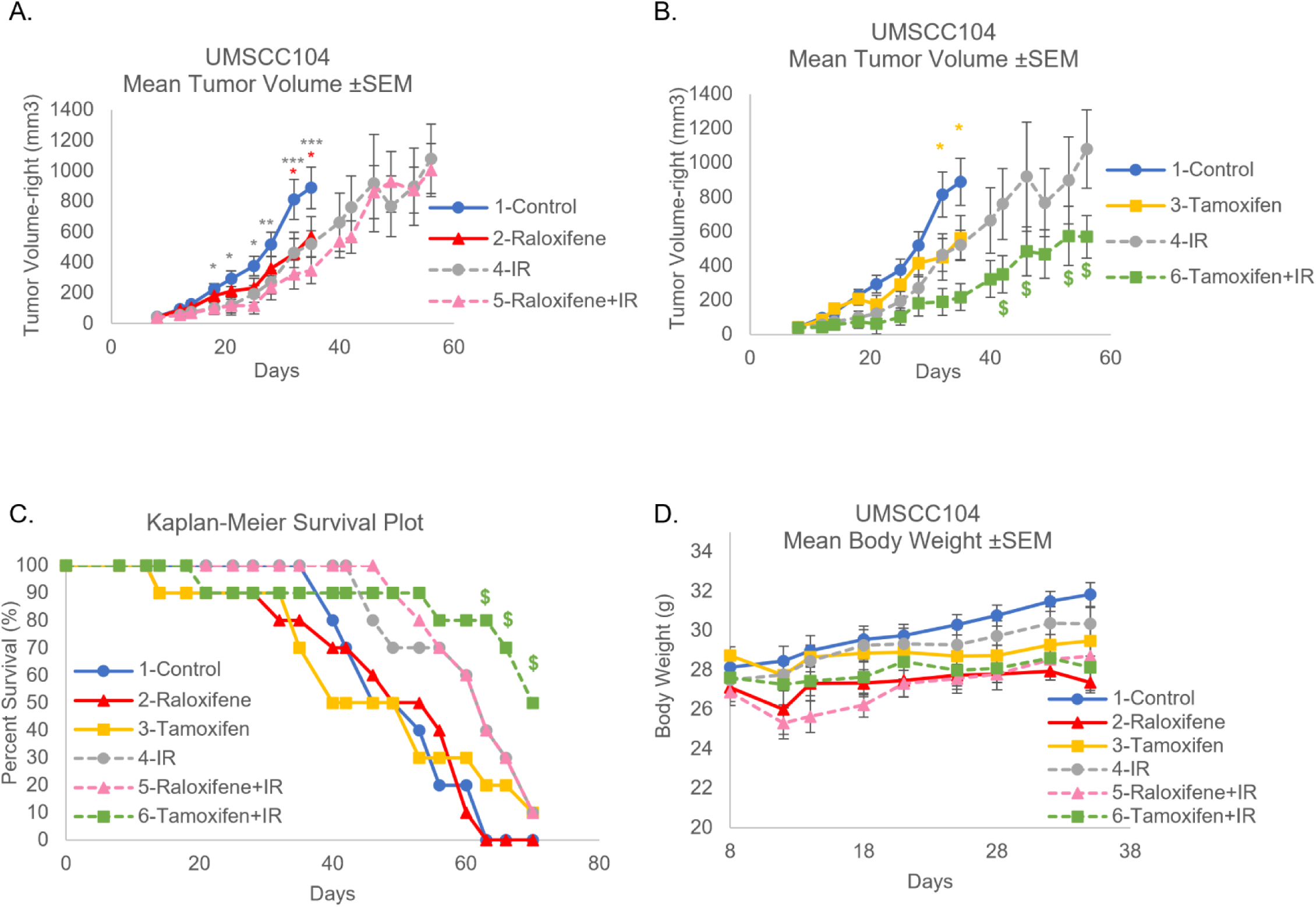
Fibroblast *in vitro* co-culture SERM assessment in UMSCC104 cells predicted the utility *in vivo*. As described in Figure 1, UMSCC104 cells were injected for xenografts in male NOD-scid IL2Rgnull (NSG) mice. Day 0 marks the date at which cells were injected for xenografts. (A,B) Tumors were palpable on day 7. 1-Control, 2-Raloxifene alone (1.5 mg), 3-Tamoxifen alone (1.0 mg), 4-radiation alone (4 Gy) (IR), as well as the combinational approaches (5-Raloxifene+IR, 6-Tamoxifen+IR) were monitored for effects on tumor volume by calipers. These experiments were run concurrently but presented on separate graphs for visual clarity (groups have been numbered for this clarity). Data points shown for Mean Tumor Volume are representative of conditions with at least 70% of animals remaining. (C) Kaplan-Meier Survival plots the survival curve of the animals treated. This data includes any mice that needed to be sacrificed due to tumor ulceration. (D) Mice were monitored for weight throughout the study. *p<0.05, **p<0.01 ***p<0.001 from control, $p<0.05 from IR alone, colors of * or $ are matched to colors of corresponding conditions that are significant from control or radiation alone, respectively.

## Discussion

We previously established that estrogen attenuates the growth of HPV^+^ keratinocyte and cancer cell lines in both an LCR and E6E7 dependent manner^25,26^. Of note, when these studies were conducted, we did not supplement fibroblast support during drug treatment^26^. Conversely, when estrogen was utilized to treat HPV^+^ xenografts in NSG mice, responsiveness was no longer observed (Figure 1). This data is supportive of previous observations by the Lambert laboratory^27^. We acknowledge the importance of the immune system in the context of HPV, the responsiveness to estrogen, and its potential impact on translational approaches as well^27,28,45,113^. As NSG mice possess a significantly compromised immune system, we then sought to investigate the role that stromal support may play in the altered responsiveness to estrogen when moving from *in vitro* to *in vivo* models. In doing so, we found that there is an HPV-specific change in response to estrogen when comparing co-culture to non-fibroblast conditions (Figure 2,3,4). We find that this response is, at least in part, HPV16 LCR dependent; however, there are likely other mechanisms at play (Figure 4). Additional mechanisms behind the observed stromal growth alterations, as well as the response to therapeutics, are currently under investigation in our laboratory and will be the subject of future reports.

Of note, the analysis of alternative species fibroblasts and mesenchymal cell types on basement membrane arrangement and growth characteristics of organotypic raft cultures were conducted many years ago^114^. Tissue-specific, species-specific, and spatial-specific alterations were found to impact numerous epithelial phenotypes^114^. Extracellular matrix components were found to have the greatest impact in this publication. We propose that the differences in extracellular matrix components between mouse and human might have moderate plating efficiency or growth efficiency alterations, and contribute to the altered growth potential observed in HeLa cells (Figure 3,4). As seen in Figure 3, UMSCC47 cell growth was not impacted by J2s; alternatively, UMSCC104 and HN30 cells grew better in the presence of J2. Regardless of the altered growth potential of cell lines in the presence or absence of fibroblasts, the observation that estrogenic sensitivity was lost in co-culture remains the same.

Anecdotally, is it known in the HPV field that J2s are crucial to the culture of primary keratinocytes for the maintenance of viral genome stability^75,76,104^. Expanding the accepted fibroblast co-culture system for HPV^+^ primary keratinocytes, our data demonstrates that this model promotes translational utility. When keratinocytes or cancer cells were grown in this co-culture system, *in vivo* results were more predictable. These models allowed for us to translate our estrogen results into estrogen pathway targeting drugs. Previous studies have highlighted the potential utility of SERMs in HPV^+^ cervical cancers, most specifically the use of raloxifene to reduce neoplastic recurrance^45,46,48,111^. We have expanded SERM analysis from cervical cancer to oropharyngeal cancer. We now demonstrate that SERMs may present clinical applicability as adjuvant approaches in HPV^+^OPC and future investigations are warranted. While our observations apply both to primary keratinocytes and cancer models of HPV, we are currently in collaboration to expand analysis to other cancer treatment models. Nevertheless, stroma is recognized to contribute to the response to therapeutics and the development of resistance, and we conclude that it should be more often considered in future translational approaches^33,35,39,40,92^.

## Materials and Methods

### Cell Culture

HN30 (generous gift from Hisashi Harada, VCU Philips Institute), UMSCC47 (Millipore; Burlington, MA, USA), and HeLa (generous gift from Alison McBride, NIAID) cells were grown in Dulbecco’s modified Eagle’s medium (DMEM) (Invitrogen; Carlsbad, CA, USA) supplemented with 10% charcoal/dextran stripped fetal bovine serum (Gemini Bio-products; West Sacramento, CA, USA). UMSCC104 (Millipore) cells were grown in Eagle’s minimum essential medium (EMEM) (Invitrogen) supplemented with nonessential amino acids (NEAA) (Gibco) and 20% charcoal/dextran stripped fetal bovine serum. N/Tert-1 cells and all derived cell lines have been described previously and were maintained in keratinocyte-serum free medium (K-SFM; Invitrogen), supplemented with a 1% (vol/vol) PenStrep (Gibco) and previously described antibiotics^20–22,115–120^. HFK^+^HPV16 have been previously described and were grown in Dermalife-K complete media (Lifeline Technology), and maintained on inactivated fibroblast feeder cells (described below)^121^. HFK+E6/E7 were grown in Dermalife-K complete media (Lifeline Technology), and maintained on inactivated fibroblast feeder cells (described below); the immortalization process is described below. Of note, we have no issues with fibroblast plating efficiency, or keratinocyte viral episome maintenance utilizing these keratinocyte complete media kits over the traditional use of E-media^121,122^. For all cells not directly purchased from companies, the cell type was confirmed by Johns Hopkins or MD Anderson cell line authentication services, and the cells were maintained at 37°C in a 5% CO_2_–95% air atmosphere, passaged every 3 or 4 days, and routinely monitored for mycoplasma (Sigma, MP0035).

### Flank xenografts for *in vivo* drug trials

HeLa and UMSCC104 cells were stably transduced with a lentiviral vector for pLX304 Luciferase-V5 blast (generous gift from Renfeng Li, originally obtained from Kevin Janes, Addgene plasmid # 98580)^123^. Cell lines were selected with 10µg/ml blasticidin. Expression was verified with bioluminescent imaging, further outlined and defined in transcriptional activity analysis method detailed below.

Xenografting was performed using previously described methodology, in collaboration with the Virginia Commonwealth University (VCU) Cancer Mouse Models Core Laboratory (CMMC)^124,125^. All experiments were conducted in accordance with animal protocol AD10002330 approved by VCU Institutional Animal Care and Use Committee. NOD-SCID-IL2γ receptor null (NSG) mice (6-8 weeks old) were injected with 1x10^6^ cells suspended in PBS and Cultrex™ basement membrane extracts (BME) (Bio-techne/R&D Systems) into the right flank. HeLa studies were conducted in female mice; UMSCC104 studies were conducted in male mice (chosen to mimic sex from human donors). Days 1-3 post-xenograft and at varying times throughout the studies, bioluminescence imaging was performed using a Xenogen IVIS-100 system (Calipers Life Sciences, Hopkinton, MA) to verify xenograft establishment, growth, and possible metastasis based on previously established protocols^126^. Tumor volume was also measured on noted dates and calculated as V =AB2 (π/6), where A is the longest dimension of the tumor, and B is the dimension of the tumor perpendicular to A. Data points presented on the graph are representative for each condition while more than 70% of the animals remained in each group. HeLa cells were palpable on day 10, UMSCC104 cells were palpable on day 7. It should be noted that in our hands, UMSCC104 xenografts are prone to ulceration. Animals were humanely euthanized when ulcerations were observed.

### NSG estrogen delivery

A combinational approach for estrogen delivery was based on modified protocols designed in collaboration with the source of our obtained control and estrogen beeswax pellets (0.4mg estrogen, Huntsman Cancer Institute, University of Utah)^127,128^. Estrogen was also delivered in drinking water using a protocol kindly shared online by the Wicha lab^129^. Briefly, a 2.7 mg/mL stock of 17-estradiol (Sigma # E2758) in ethanol was diluted to a final concentration of 8 µg/mL in sterile drinking water. Pellets (control or estrogen) were implanted after tumors were palpable. Drinking water supplementation also began at this time point. “Early estrogen” in UMSCC104 denotes that water estrogen supplementation began the same day as xenograft injections (pellets were still implanted when tumors became palpable on day 7).

### NSG SERM delivery

Treatment of mice with Raloxifene was performed as previously described by the Lambert Laboratory^45,111^. The human formulation of raloxifene hydrochloride (60mg tablets; EVISTA; Eli Lilly) were purchased from Virginia Commonwealth University Health System Pharmacy. Tablets were resuspended in PBS for a final concentration of 10 mg/ml. Mice were administered a 150 μl drug suspension (equivalent to 1.5 mg) via i.p. injection. Treatment of Tamoxifen was performed as previously described^130^. Tamoxifen (Sigma, T-5648) was resuspended in corn oil (Sigma C-8267) at 37°C for a final concentration of 10 mg/ml. Mice were administered 100 µl drug suspension (equivalent to 1.0 mg) via i.p. injection. SERM treatments began after tumors were palpable. Mice received treatment 5 days a week for 4 weeks, for a total of 20 injections.

### Small Animal Radiation Research Platform (SARRP) ionizing radiation delivery

1 day following pellet implantation to allow for mouse recovery, targeted ionizing radiation (IR) was delivered utilizing the Xstrahal SARRP. HeLa studies utilized 10 Gy. Due to the radiation sensitivity of NSG mice, UMSCC104 studies reduced IR dose to 5 Gy, and finally to 4 Gy.

### Culture, conditioned media collection, and mitomycin C (MMC) inactivation of 3T3-J2 mouse feeder cells

3T3-J2 immortalized mouse fibroblasts (J2) were grown in DMEM and supplemented with 10% FBS. Fresh media was exchanged twice a week; conditioned media was spun down as 500 rcf to remove any residual cells^131,132^. 80-90% confluent plates were supplemented with 4µg/ml of MMC in DMSO (Cell Signaling Technology) for 4-6 hours at 37°C. MMC-supplemented medium was removed and cells were washed with 1xPBS. Cells were trypsinized, spun down at 500 rcf for 5 mins, washed once with 1xPBS, spun again, and resuspended. Quality control of inactivation (lack of proliferation) was monitored for each new batch of mitomycin-C. 100-mm plate conditions were supplemented with 1x10^6^ J2 and 6-well plate conditions were supplemented with 1x10^5^ every 2-3 days; for longer term cultures, any remaining J2s were washed off with 1x PBS and new J2 were continually supplemented every 2-3 days.

### Culture and mitomycin C (MMC) inactivation of human dermal mesenchymal fibroblast feeder cells

Human dermal mesenchymal fibroblasts (HDFM) were grown, treated, and quality controlled as described for the above J2 protocol. 6-well plate conditions were supplemented with 1x10^5^ every 2-3 days.

### Generation of E6E7-immortalized keratinocytes

Primary keratinocytes from single donors were obtained from LifeLine Cell Technologies^121^. Cells were cultured on collagen-coated plates for lentiviral delivery of HPV16 E6E7, using the pLXSN16E6E7 plasmid (Addgene plasmid # 52394, a gift from Denise Galloway)^133^. Following selection with G418 (72mM), cells were cultured on mitomycin-C inactivated 3T3-J2 fibroblasts. HFKs were cultured in DermaLife-K Complete media (LifeLine Cell Technologies) and E6E7 expression was confirmed by qRTPCR. Of note, these cells were generated at the same time and utilizing the same donors as the previously described HFK-HPV16^121^.

### Co-culture of keratinocytes in the presence or absence of inactivated fibroblasts

N/Tert-1, HFK E6E7, or HFK HPV16 cells were seeded at 5x10^5^ in 100-mm plates in the presence or absence of previously seeded J2s (at least 6 hours prior). Twenty-four hours later, noted cells were supplemented with 15 μM 17β-estradiol (estrogen). Forty-eight hours after estrogen supplementation, plates were washed to remove residual J2 and cells were trypsinized and counted. For analysis of 1 week time point, HFK E6E7 or HFK HPV16 cells were seeded at 1x10^5^ in 100-mm plates in the presence or absence of previously seeded J2s. J2s were re-supplemented every 2-3 days as previously described. Pellets from these experiments were utilized for subsequent immunoblotting or DNA analysis, detailed below.

### Generation of stable nuclear labeled cells with Incucyte® Nuclight Lentivirus

mKate2 Incucyte® Nuclight Lentivirus (puro) cells were generated according to the Sartorius product guide protocol (Sartorius cat# 4476), using a MOI of 3 or 6, depending on cell type. Cells generated were maintained in 1µg/ml puromycin supplemented media. Fluorescence was routinely monitored by BZ-X TexasRed filter via the Keyence BZ-X800 inverted fluorescence microscope.

### Co-culture of nuclear-labeled cancer cell lines in the presence or absence of inactivated fibroblasts

Stable mKate2-puro HeLa, UMSCC47, UMSCC104, or HN30 cells were seeded in triplicate at 1x10^4^ per well in 6-well plates in the presence or absence of previously seeded J2 cells, or HDFM cells (at least 6 hours prior – fibroblast type is noted for each experiment). Twenty-four hours after seeding, day 0 images were captured in brightfield and TexasRed with the Keyence BZ-X800 Image Cytometer. Noted drugs were supplemented immediately after this initial imaging: 1.5µM (HeLa) or 15 μM 17β-estradiol (Sigma), 10µM Tamoxifen (MP Biomedicals), 10µM Raloxifene (Cayman Chemical Company), or 10-20µM Cisplatin (APExBIO). Cytometry images were again captured on day 1 and day 3. UMSCC104 were washed on day 3 after imaging, new J2 and drug were supplemented, and additional images were captured on day 5 and 7. Ten fields of view were randomized per well for all conditions. Cell count image cytometry batch analysis was performed using the Keyence BZ-X800 Image Analyzer software. All conditions utilized set analysis conditions from a single randomized control image and applied to all data points automatically to reduce variability and bias. Data is presented as fold of control from day 0. Representative images are presented in Figure 9.

**Figure 9.**
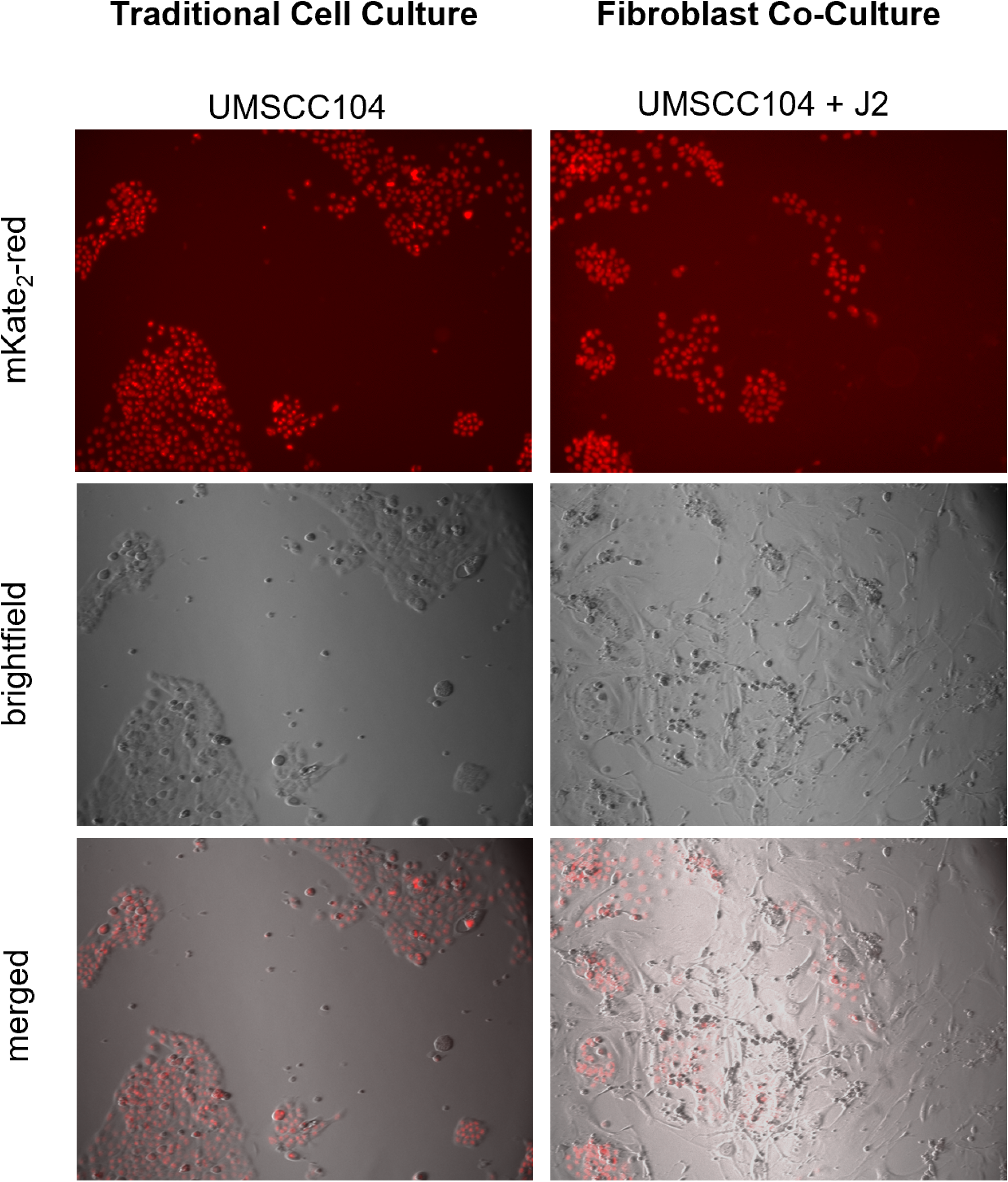
Representative cell culture images quantified via Keyence. Images presented are day 5 of UMSCC104 cell cultures in the presence or absence of J2s. Images were captured at 10x in brightfield and TexasRed. All cell lines at all time points were counted utilizing the same control-established automated parameters to ensure reproducibility.

### Immunoblotting

Specified cells were washed with 1X PBS and trypsinized. Pellets were washed with 1X PBS and resuspended in 5X packed cell volume of NP40 buffer (50mM Tris-HCl Ph 7.5, 150mM NaCl, 1%NP-40, 5mM EDTA) supplemented with Roche cOmplete protease inhibitor and Roche PhosSTOP phosphatase inhibitor. Cell-lysis buffer suspension was incubated on ice for 30 min with occasional agitation, then centrifuged for 15 min at 14,000 rcf at 4 °C. Supernatant protein concentration was measured via the Bio-Rad protein estimation assay according to manufacturer’s instructions. 100 μg protein samples were heated at 95 °C in 4x Laemmli sample buffer (Bio-Rad) for 5 min. Noted samples were run down a Novex 4–12% Tris-glycine gel (Invitrogen) and transferred onto a nitrocellulose membrane (Bio-Rad) at 30V overnight, or 100V for 1 hour using the wet-blot transfer method. Membranes were blocked with Odyssey (PBS) blocking buffer (diluted 1:1 with 1X PBS) at room temperature for 1 hour and probed with indicated primary antibody diluted in Odyssey blocking buffer. Membranes were washed with PBS supplemented with 0.1% Tween (PBS-Tween) and probed with the indicated Odyssey secondary antibody 1:10,000 (goat anti-mouse IRdye 800CW or goat anti-rabbit IRdye 680CW) diluted in Odyssey blocking buffer. Membranes were washed three times with PBS-Tween and an additional wash with 1X PBS. Infrared imaging of the blot was performed using the Odyssey CLx Li-Cor imaging system. Immunoblots were quantified using ImageJ utilizing GAPDH as the internal loading control. The following primary antibodies were used for immunoblotting in this study at 1:1000, unless otherwise noted: ERα (Abcam, ab32063), GAPDH 1:10,000 (Santa Cruz, sc-47724), pRb (Santa Cruz, sc-102), p53 (Cell Signaling Technology, CST-2527 and CST-1C12), γH2AX 1:500 (Cell Signaling Technology, CST-80312 and CST-20E3), SIRT1 (EMD Millipore 07-131), β-actin (Santa Cruz, sc-47778), E7 1:500 (Santa Cruz, sc-6981).

### Transfection and transcriptional activity analysis

N/Tert-1 cells were plated at a density of 5 × 10^5^ in 100-mm dishes. The following day, the previously described plasmids for pGL3 basic, pGL3 control, or pHPV16-LCR-Luc were transfected Lipofectamine 2000 (according to the manufacturer’s instructions, ThermoFisher Scientific). Twenty-four hours after transfection, cells were washed, and noted cells were supplemented with 15 μM 17β-estradiol; J2s were also supplemented at this time point for noted conditions to reduce likelihood of altered transfection efficiency. Forty-eight hours after transfection, cells were harvested utilizing Promega reporter lysis buffer and analyzed for luciferase using the Promega luciferase assay system. Concentrations were normalized to protein levels, as measured by the Bio-Rad protein assay dye. Relative fluorescence units (RFU) were measured using the BioTek Synergy H1 hybrid reader.

### Exonuclease V assay

PCR based analysis of viral genome status was performed using methods described by Myers *et al.* ^84^. Briefly, 20 ng genomic DNA was either treated with exonuclease V (RecBCD, NEB), in a total volume of 30 ul, or left untreated for 1 hour at 37°C followed by heat inactivation at 95°C for 10 minutes. 2 ng of digested/undigested DNA was then quantified by real time PCR using a 7500 FAST Applied Biosystems thermocycler with SYBR Green PCR Master Mix (Applied Biosystems) and 100 nM of primer in a 20 μl reaction. Nuclease free water was used in place of the template for a negative control. The following cycling conditions were used: 50°C for 2 minutes, 95°C for 10 minutes, 40 cycles at 95°C for 15 seconds, and a dissociation stage of 95°C for 15 seconds, 60°C for 1 minute, 95°C for 15 seconds, and 60°C for 15 seconds. Separate PCR reactions were performed to amplify HPV16 E6 F: 5’- TTGCTTTTCGGGATTTATGC-3’ R: 5’- CAGGACACAGTGGCTTTTGA-3’, HPV16 E2 F: 5’- TGGAAGTGCAGTTTGATGGA-3’ R: 5’- CCGCATGAACTTCCCATACT-3’, human mitochondrial DNA F: 5’-CAGGAGTAGGAGAGAGGGAGGTAAG-3’ R: 5’- TACCCATCATAATCGGAGGCTTTGG -3’, and human GAPDH DNA F: 5’- GGAGCGAGATCCCTCCAAAAT-3’ R: 5’- GGCTGTTGTCATACTTCTCATGG-3’

### Reproducibility, research integrity, and statistical analysis

All *in vitro* experiments were carried out at least in triplicate in all of the cell lines indicated. All cell lines were bought directly from sources indicated, or typed via cell line authentication services. All images shown are representatives from triplicate experiments. *In vivo* experiments were designed in collaboration with the VCU Massey Cancer Center animal core and biostats core for sample size justification and statistical power analysis. Quantification is presented as mean +/- standard error (SE). Student’s t-test or analysis of variance (ANOVA) were used to determine significance as appropriate: * p<0.05, **p<0.01, ***p<0.001

## Data availability

Following the 2023 NIH data management and sharing policy, all data resulting from the development of projects will be available in scientific communications presented at conferences and in manuscripts that will be published in peer-reviewed scientific journals. Data will be deposited in the Open Science Framework (OSF) platform. OSF can be accessed at https://osf.io. VCU is an OSF institutional member, and OSF is an approved generalist repository for the 2023 NIH data management and sharing policy.

## Acknowledgements

This work was supported by the VCU Philips Institute for Oral Health Research, the National Institute of Dental and Craniofacial Research/NIH/DHHS R03 DE029548, and the National Cancer Institute-designated Massey Cancer Center grant P30 CA016059. Mouse services and products in support of the research project were generated by the Virginia Commonwealth University Cancer Mouse Models Core Laboratory, supported, in part, with funding from NIH-NCI Cancer Center Support Grant P30 CA016059.

